# Human-specific retrotransposons encode the regulatory logic of two lineages in a bipotent neural crest and central nervous system precursor

**DOI:** 10.64898/2026.09.16.752192

**Authors:** Cong Minh Duc Nguyen, Zoe H. Mitchell, Marco Trizzino

## Abstract

Transposable elements are sources of cis-regulatory innovation, and the youngest families are particularly interesting because only recent insertions can account for species-specific regulatory divergence. SVAs (subfamilies -E, F), LTR5Hs and L1HS are human-specific retroelements. They are polymorphic, and in some cases still mobile. Regulatory activity in these families has been surveyed mainly in pluripotent cells and differentiated tissues, leaving the early embryonic precursors in which lineage decisions are made largely unexplored. We profiled chromatin accessibility and gene expression in human iPSC-derived bipotent neuroectodermal aggregates harvested at a stage that precedes lineage commitment and retains competence for both the neural crest and the central nervous system. Transcriptional profiling confirmed this dual competence: neural plate border, neural crest and forebrain programs were concurrently active, and definitive neural crest specifiers not yet induced.

Of ∼4,300 annotated human-specific transposons, 289 were reproducibly accessible in the aggregates. Relative to inaccessible elements, these were closer to genes, occupied gene-dense neighborhoods, and had more highly expressed neighboring transcripts. They were enriched for transcription factor binding motifs drawn from both neural crest and neuronal programs, mirroring the dual competence of the cells. Co-option was strongly family-biased in both extent and content, with SVA-F and LTR5Hs over-represented among accessible elements and each family carrying a distinct motif repertoire. Interestingly, protein interaction analysis of the genes near accessible elements recovered a module built around TRIM28 and multiple KRAB zinc-finger proteins, the machinery that silences transposons through H3K9me3 suggesting a potential feedback loop. These data identify human-specific elements as candidate regulatory substrates at a bipotent developmental stage.

## Introduction

Transposable elements (TEs) constitute roughly half of the human genome and have historically been viewed mainly as source of genomic instability, capable of generating insertional mutations, chromosomal rearrangements and a range of monogenic and complex diseases. However, over the past two decades, a substantial body of evidence has established that TEs, including human-specific ones, are also repeatedly co-opted by the host genome as functional cis-regulatory elements (CREs), including enhancers, alternative promoters, insulators and chromatin domain boundaries (Wang et al. 2007; Bourque et al. 2008; Sasaki et al. 2008; Markljung et al. 2009; Kunarso et al. 2010; Lynch et al. 2011, 2015; Jacques et al. 2013; Chuong et al. 2013, 2016; Sundaram et al. 2014; Trizzino et al. 2017, 2018; Pontis et al. 2019; Barnada et al. 2022; Patoori et al. 2022; Fueyo et al., 2022, 2025; Deelen et al. 2025). Co-option most commonly proceeds through the accumulation of mutations within an ancestral TE sequence that fortuitously create or strengthen a binding site for host transcription factors. Because this process depends on the pre-existing sequence composition of each TE family, different TE families show markedly different propensities to be co-opted by different transcription factors (Bourque et al., 2008; Schmidt et al. 2012).

TE activity, and hence the opportunity for co-option, is markedly elevated during gametogenesis and early embryonic development, reflecting the evolutionary pressure on TEs to propagate within the germline before epigenetic silencing is re-established. In human pluripotent cells, TE-derived elements function as binding platforms for core pluripotency transcription factors including OCT4, SOX2 and NANOG (Kunarso et al., 2010; Barnada et al. 2022), and TE-derived enhancers have been directly implicated in placental gene regulation and immune response (Lynch et al. 2015; Frost et al. 2023; Chuong et al., 2013). Because early embryonic and extra-embryonic gene regulatory networks are amongst the most rapidly evolving in mammalian genomes, and since TE insertion is one of the principal mechanisms by which entirely new regulatory elements can arise over short evolutionary timescales, TE co-option is now considered a major contributor to species-specific, and in particular human-specific, gene regulatory innovation (Trizzino et al., 2017, 2018, Fueyo et al. 2025).

Neural crest cells (NCCs) arise from the neural plate border, a transient territory of the dorsal ectoderm positioned at the interface between the prospective neural plate (which will form the Central Nervous System [CNS]) and the non-neural, prospective epidermal ectoderm. As the neural plate folds and the neural tube closes, neural plate border cells converge dorsally and, under the combinatorial influence of multiple signaling gradients, diverge into at least three distinct derivatives: the neural crest, the cranial sensory placodes, and the dorsal CNS neuroepithelium (Mayor & Theveneau, 2013; Bronner & Simões-Costa, 2016). Critically, the neural plate border gene regulatory network is built from a shared, overlapping set of transcription factors, including ZIC, PAX3/7, MSX and TFAP2 family members, that are initially co-expressed across the prospective neural crest, placodal and dorsal CNS territories, with fate becoming progressively restricted as development moves forward (Simões-Costa & Bronner, 2015). Single-cell transcriptomic studies of pre-migratory and early migratory neural crest have confirmed that this population is transcriptionally heterogeneous even prior to lineage commitment, with subsets of cells co-expressing markers more typically associated with differentiated CNS or sensory neuronal derivatives well before migration or terminal differentiation has occurred (Simões-Costa et al., 2014; Lumb et al., 2017; Lencer et al., 2021; Keuls et al., 2023; Demurtas et al. 2025).

This developmental framework has direct relevance to the interpretation of *in vitro* NCC differentiation models. The commonly used three-dimensional based protocol (Bajpai et al. 2010; Prescott et al 2015; Ozga et al. 2026) does not differentiate iPSCs directly into a homogeneous, already-fate-restricted NCC population. Rather, it generates a neuroectodermal, neural-plate-border-like precursor population (hereafter neuroectodermal aggregates) that is competent to generate both neural crest and CNS/sensory neuronal derivatives, mirroring the *in vivo* bipotency of this lineage. Notably, this protocol has been widely used to generate and study migratory cranial neural crest at the endpoint of the protocol (∼day 14; Prescott et al 2015; Long et al. 2020; Demurtas et al 2025; Mitchell et al. 2025). However, the bipotent, transient pre-migratory state it generates (neuroectodermal aggregates) has remained largely unprofiled.

Despite extensive genomic similarity between humans and other great apes, craniofacial morphology and brain development diverge markedly across Hominidae. Human-specific TEs (predominantly L1HS, LTR5Hs, and SVA) domesticated as cis-regulatory elements represent the subset of the repeat landscape that is both young enough to underlie this divergence and variable enough to contribute to phenotypic variation within the species. Work from our group has shown that the human-specific retroelement family LTR5Hs is required for late neural crest migration (Deelen et al., 2025), and that TE families exclusive to the human lineage (SVAs) drive distinct gene regulatory programs in human pluripotent stem cells and in hippocampal intermediate progenitor cells (Barnada et al., 2022; Patoori et al., 2022). SVAs are of particular interest: the human-specific SVA-E and SVA-F subtypes first colonized our lineage roughly 3 million years ago, are present in almost 3000 annotated copies, are still mobile and are highly polymorphic (Wang et al., 2005; Hancks & Kazazian, 2010). The regulatory impact of these families is disproportionate to their abundance: SVAs account for roughly one-thousandth of the genomic TE content, yet they represent 17% of TE-derived sequences bearing active enhancer marks in fetal brain (Pontis et al., 2019).

Human-specific elements are therefore likely to have contributed to the regulatory divergence underlying both craniofacial and neural evolution in our lineage. However, despite the evidence for TE co-option in migratory NCCs, pluripotent cells and CNS progenitors, almost nothing is known about the pre-migratory, neural-plate-border-equivalent stage that gives rise to both lineages, even though it establishes the regulatory landscape on which later NCC migration and CNS specification both depend. (Simões-Costa & Bronner, 2015; Simões-Costa et al., 2014). In this framework, we mapped genome-wide chromatin accessibility and gene expression in human iPSC-derived neuroectodermal aggregates and identified human-specific TEs with candidate cis-regulatory activity. We characterized the genomic distribution, motif content and TE-family-specific co-option biases of these accessible elements and examined their contribution to developmental gene expression.

## Results

### An iPSC-derived model of the bipotent NCC/CNS-competent neuroectoderm

We generated pre-migratory, NCC/CNS-competent neuroectodermal aggregates from human iPSCs (Bajpai et al. 2010; **Fig. 1A**). This protocol was selected specifically because it recapitulates neural-plate-border-like bipotency *in vitro*, generating neuroectodermal aggregates competent to give rise to both neural crest and CNS/sensory neuronal derivatives, rather than a pre-restricted, NCC-only population. By day 5, the collection timepoint for this study, the aggregates displayed the compacted, stratified cellular arrangement characteristic of neuroectodermal identity. Critically, this timepoint precedes neural crest cell specification and migration, which usually happens in culture several days later (Bajpai et al. 2010, Prescott et al. 2015; Ozga et al. 2026).

**Figure 1.**
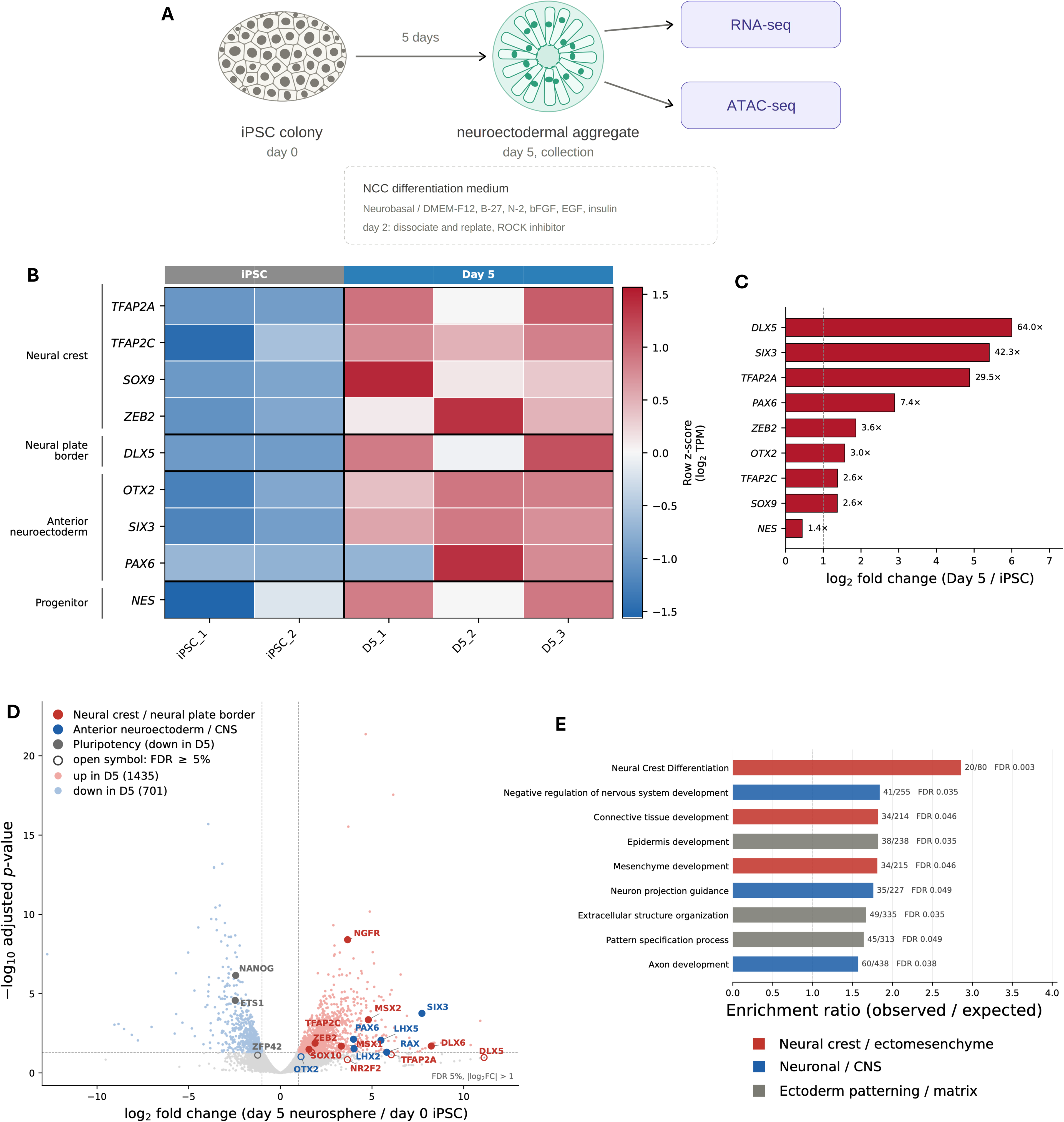
Day-5 neuroectodermal aggregates acquire a bipotent neuroectodermal identity. (A) Differentiation scheme. iPSC colonies were differentiated for 5 days in NCC medium and collected for parallel RNA-seq and ATAC-seq. (B) Expression of lineage markers by RNA-seq (iPSC, day-5). Color shows row z-score of log2 TPM. Genes are grouped by functional category. (C) Log2 fold change (day 5 / iPSC) for the same markers, with linear fold change annotated. Dashed line marks 2-fold. (D) Volcano plot of differential expression (day 5 vs iPSC). Colored points pass FDR < 5% and |log2FC| > 1; selected lineage markers are labelled, with open symbols indicating FDR ≥ 5%. (E) Over-representation analysis of upregulated genes. Bars show enrichment ratio (observed/expected). Overlap counts and FDR are annotated. Colors group terms by biological theme.

Identity was confirmed by RNA-seq, performed in iPSCs and at day-5, which showed that the aggregates comprise an uncommitted anterior neuroectodermal population in which neural crest specification (high *SOX9*, *TFAP2A*, *ZEB2*; **Figs. 1B, C**), neural plate border (*DLX5*) and forebrain-territory identity (*SIX3, OTX2, PAX6*; **Figs. 1B, C**) are concurrently active, with cells having exited pluripotency but not yet acquired migratory neural crest (e.g. *SOX10* not yet expressed; Supplementary File 1) or definitive neuroepithelial character (*SOX1*-negative suggests no dorsoventral patterning).

To further examine the cell identity of the neuroectodermal aggregates, the full transcriptome of iPSCs and day-5 cells was analyzed using DESeq2 (Love et al. 2014). Differentiation produced a substantial transcriptional shift: 2,136 genes were differentially expressed between the two time points (FDR < 0.05; | log2FC | > 1), of which 1,435 were upregulated and 701 downregulated at day 5 (**Fig. 1D**; Supplementary File 2). The genes most strongly induced at day 5 support and identify the population as neural plate border-derived and anteriorly patterned. *SIX3, LHX5, RAX, LHX2* and *PAX6* were all induced essentially from zero, establishing an anterior neural character. Consistent with this, *OTX2* was the most abundant of the markers examined. In parallel, the neural plate border and neural crest arm of the program was induced: *MSX1/2, DLX5/6, TFAP2A/C, ZEB2, NGFR* were all significantly upregulated (**Fig. 1D**; Supplementary File 2). *SOX9* was not significantly upregulated and *SOX10* not expressed, placing the population at the onset of neural crest specification rather than at an established stage, consistent with the temporal order in which the AP-2 coordinators precede SOX9/10.

To investigate which biological programs were enriched in the list of induced (upregulated) genes, we performed over-representation analysis on the upregulated genes using WebGestalt. The most strongly enriched set in the analysis was Neural Crest Differentiation (FDR 2.9 × 10⁻^3^; **Fig. 1E**; Supplementary File 3), providing an unbiased confirmation of neural crest identity that is independent of the marker genes examined above. Neural crest derivative programs were enriched alongside it, including mesenchyme and connective tissue development (**Fig. 1E**; Supplementary File 3), consistent with the ectomesenchymal potential of cranial neural crest. Critically, neuronal and central nervous system programs, including axon development and neuron projection guidance were also enriched in the same, unstratified gene list (**Fig. 1E**; Supplementary File 3). Enrichment of “epidermis development” further reflects the position of the neural plate border at the interface between neural plate and non-neural ectoderm. The co-occurrence of neural crest, neuronal and non-neural ectodermal signatures within a single population, rather than the dominance of any one of them, is the pattern predicted for a neural plate border-equivalent state in which these fates have not yet been resolved.

Together, these observations confirm that day-5 cell aggregates correspond to an early, pre-migratory neuroectodermal population appropriate for genome-wide profiling of the shared, not-yet-lineage-restricted NCC/CNS regulatory landscape.

### Accessible human-specific TEs occupy gene-dense genomic neighborhoods

ATAC-seq libraries were generated from day-5 neuroectodermal aggregates in two independent replicates (paired-end 150 bp reads; peak calling FDR < 5%). Of 4,294 annotated human-specific TEs genome-wide, 289 (6.7%; Supplementary File 4) showed reproducible accessibility across both replicates (**Fig. 2A**). We next asked whether these accessible TEs occupy genomic positions consistent with a cis-regulatory role. Indeed, accessible human-specific TEs were located significantly closer to the nearest gene TSS than non-accessible human-specific TEs (15.2 Kb vs 34.5 Kb, Wilcoxon rank-sum test, p = 3.3 × 10⁻^15^; **Fig. 2B**; Supplementary File 5). At the same time, the great majority of accessible TEs (281/289, 97.2%) lay more than 1,000 bp from the nearest TSS, a distance more consistent with a distal enhancer-like element than with a core promoter.

**Figure 2.**
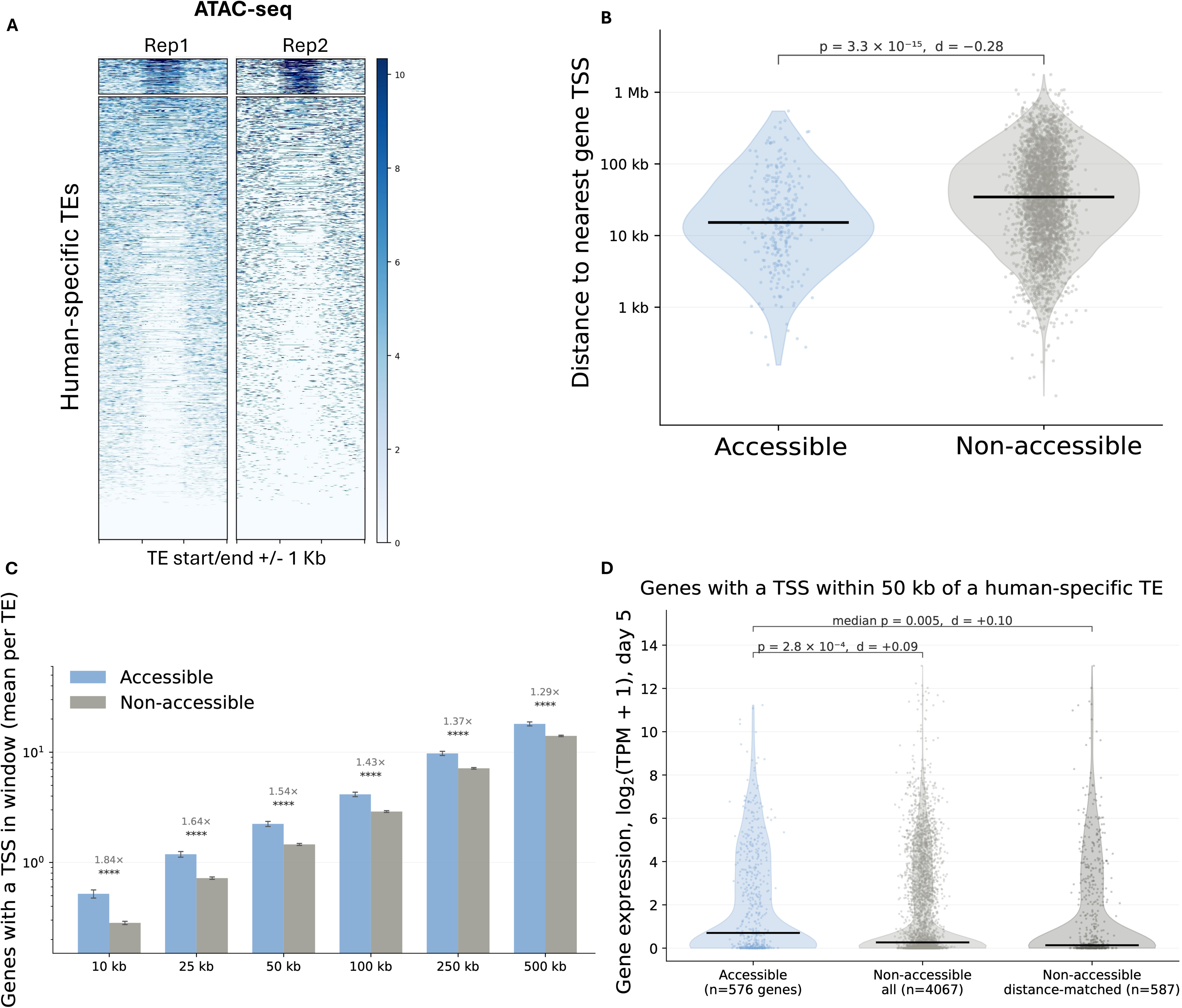
Accessible human-specific TEs are closer to genes and sit in gene-dense regions. (A) ATAC-seq signal across human-specific TEs in two day-5 replicates, ±1 kb around element boundaries, sorted by mean signal. (B) Distance to nearest gene TSS for accessible (n = 289) versus non-accessible elements. Violins show the distribution, points individual elements, bars the median. Two-sided Mann– Whitney U with Cliff’s delta. (C) Mean number of genes with a TSS within the indicated window, per element. Error bars show SEM. Fold difference and significance are annotated. ****P < 0.0001. (D) Day-5 expression of genes with a TSS within 50 kb of an accessible element, compared with all non-accessible elements and with a distance-matched control set. Bars show medians. Statistics as in (B), with the matched comparison reporting the median across resamples.

Having established that accessible human-specific TEs lie closer to transcriptional start sites than non-accessible elements, we asked whether this reflects a general difference in the genomic environment these elements occupy. For each annotated human-specific TE, we counted the genes whose TSS fell within a series of windows centered on the element (from 10 kb windows to 500 kb windows), restricting the count to the 33,588 genes quantified in our RNA-seq data. Accessible TEs were consistently embedded in more gene-dense regions than non-accessible TEs, at every scale examined (**Fig. 2C**). Within 10 kb, accessible elements had a mean of 0.52 genes per TE compared with 0.28 for non-accessible elements, a 1.84-fold difference (Mann-Whitney p = 2.7 × 10⁻^11^). The difference persisted across all larger windows, at 1.54-fold within 50 kb (2.24 vs 1.46 genes per TE, p = 4.3 × 10⁻^15^), 1.43-fold within 100 kb and 1.29-fold within 500 kb (all p < 10⁻^9^). Notably, the magnitude of the enrichment declined gradually with window size, indicating that the excess of genes is concentrated in the immediate vicinity of accessible elements rather than reflecting their assignment to broadly gene-rich chromosomal territories.

### Genes surrounding accessible TEs are more highly expressed in the neuroectodermal aggregates

We next asked whether this favorable genomic positioning is accompanied by higher transcriptional output of the surrounding genes. Considering only the single nearest gene to each element, genes adjacent to accessible TEs showed a modest trend towards higher day-5 expression that did not reach significance (median log2[TPM + 1] 0.21 vs 0.1; p = 0.066). Because a single nearest-neighbor assignment captures only a fraction of the genes a distal regulatory element might influence, and discards information from all other genes in the locus, we repeated the analysis considering every gene with a TSS within a defined window of each element.

This window-based analysis revealed a significant difference at every scale tested. Within 50 kb, the 576 genes neighboring accessible TEs were more highly expressed than the 4,067 genes neighboring non-accessible TEs (median log2[TPM + 1] 0.70 vs 0.27; p = 2.8 × 10⁻^4^; **Fig. 2D**; Supplementary File 6). The same direction of effect was observed from 10 kb (p = 9.4 × 10⁻^3^; Supplementary File 6) through to 500 kb (p = 3.3 × 10⁻^8^; Supplementary File 6), with the largest effect size at the shortest window, again pointing to a locally concentrated signal.

Since accessible TEs are positioned closer to genes than non-accessible elements, we asked whether the expression difference simply reflects this proximity. We therefore constructed control sets of non-accessible TEs matched to the accessible set on their distribution of distance to the nearest TSS, binned by distance, and repeated the comparison over 300 independent resamples of the matched control. The difference was unaffected by this correction (median Cliff’s delta +0.10, median p = 0.005), indicating that the higher expression of genes surrounding accessible TEs is not merely a consequence of their being closer to genes.

Taken together, these observations indicate that the subset of human-specific TEs that acquire chromatin accessibility in day-5 neuroectodermal aggregates is not randomly distributed with respect to the transcribed genome, as these elements are preferentially located in gene-dense regions, close to their neighboring genes, and the genes surrounding them are more transcriptionally active than those surrounding the non-accessible majority. This is the genomic context expected of elements acting as cis-regulatory sequences, although these data are correlative and do not establish that individual elements drive the expression of the nearby genes.

### Accessible TEs are enriched for transcription factor binding motifs spanning both neural crest and CNS/sensory differentiation programs

Motif analysis of the 289 accessible TE sequences identified 15 significantly enriched transcription factor binding motifs (TFBMs; E < 1 × 10⁻^5^; **Fig. 3A**; Supplementary File 7). Notably, the identity of the enriched motifs spans both neural-crest-associated and CNS/sensory-neuron-associated regulatory programs, directly mirroring the developmental bipotency of the profiled population. The single most enriched motifs corresponded to NEUROD2 and ATOH1, both of which are core drivers of neurogenesis and terminal neuronal differentiation from neuroepithelial progenitors (**Fig. 3A**). ATOH1 additionally has an essential, well-characterized role in the specification of inner ear hair cells, the mechanosensory receptor cells of the inner ear, which derive from the otic placode, a cranial sensory placode territory that is developmentally intertwined with, but distinct from, the neural crest proper (Bermingham et al. 1999; Groves and LaBonne 2014). STAT3 and PAX5 motifs were also significantly enriched (**Fig. 3A**). STAT3 acts as a switch governing the transition of neural plate border cells from an undifferentiated, high-STAT3 state to a specified, proliferative neural-crest state at lower STAT3 activity, positioning it at the neural-crest/CNS decision point (Nichane et al. 2010). PAX5, by contrast, functions in patterning the midbrain-hindbrain boundary of the neural tube and cerebellar development, a purely CNS-territory function (Urbanek et al. 1997; Hidalgo-Sanchez et al., 2022). We additionally identified motifs for CRX (**Fig. 3A**), required for photoreceptor differentiation within the CNS-derived retina (Furukawa et al. 1999; Zheng and Chen 2024), and for SMAD4 (**Fig. 3A**), which is required for NCC-derived tooth ectomesenchyme development, a clearly neural-crest-restricted function (Ko et al. 2007; Li et al. 2011).

**Figure 3.**
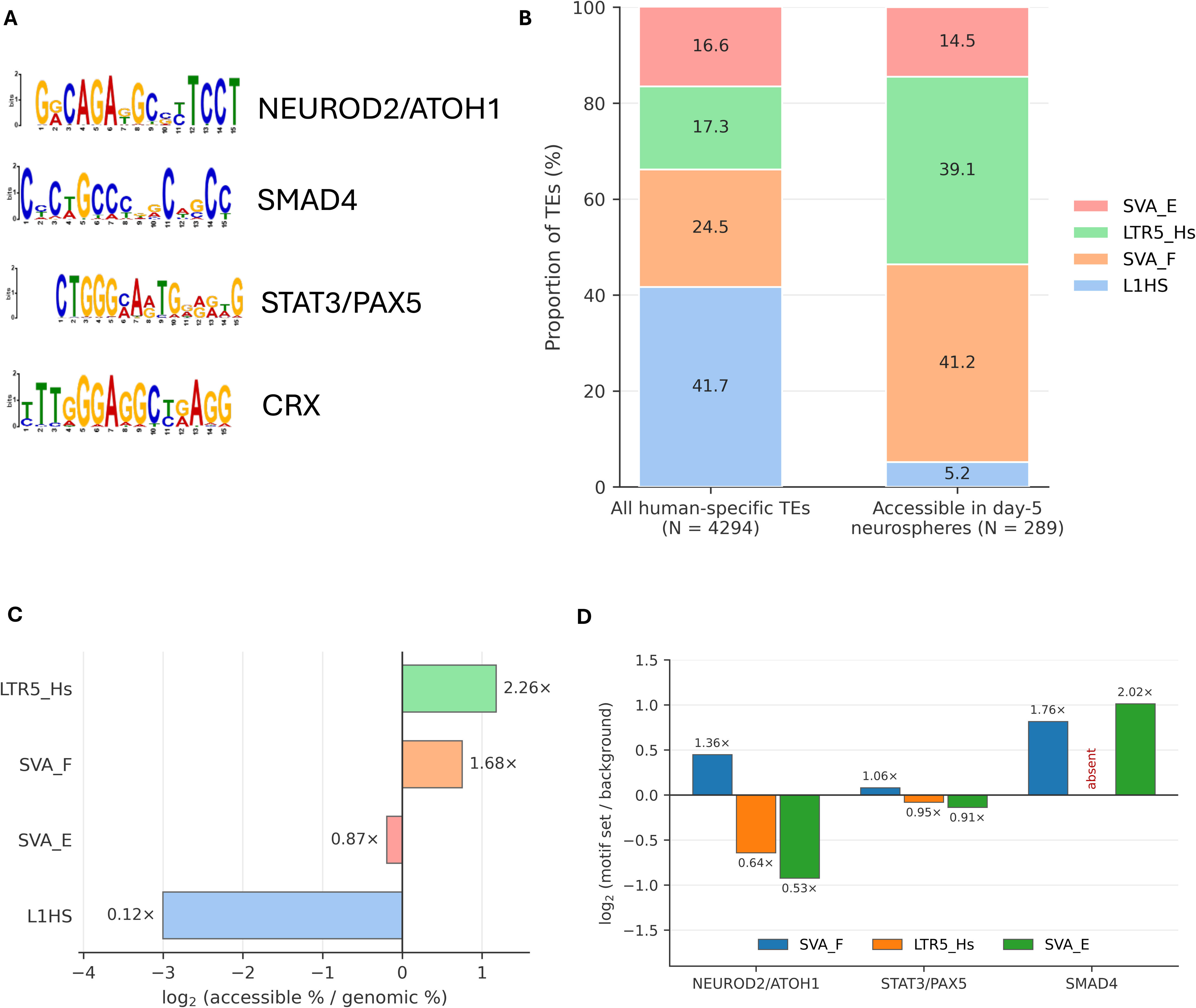
Accessibility and motif content are biased by TE family. (A) Sequence logos for the transcription factor binding motifs enriched in accessible human-specific TEs. (B) Family composition of all human-specific TEs and of those accessible in day-5 aggregates. (C) Enrichment of each family among accessible elements relative to genomic abundance. Linear fold change is annotated. (D) Family composition of each motif set relative to all motif-containing TEs. SMAD4 motifs were absent from LTR5Hs.

Overall, the co-occurrence, within the same accessible-TE motif set, of transcription factors governing terminal CNS/sensory neuronal differentiation (NEUROD2/ATOH1, CRX), the neural-crest/CNS decision itself (STAT3), CNS patterning (PAX5) and neural-crest-restricted mesenchymal differentiation (SMAD4) provides direct molecular support for the view that the accessible chromatin landscape of day-5 aggregates reflects a bipotent regulatory state rather than a committed neural crest or CNS program.

Reverse motif mapping performed with MEME-FIMO showed that 210 of 289 accessible TEs (73%) contained at least one enriched TFBM, and 172 (60%) contained more than one, with the single most motif-dense element (SVA_F_r1036) harboring 22 individual TFBM instances spanning four distinct motifs (NEUROD2/ATOH1, STAT3/PAX5 and SMAD4). The frequent co-occurrence of multiple distinct TFBMs within individual TEs suggests that many of these elements function as compact cis-regulatory modules capable of integrating several transcriptional inputs, potentially including inputs from both the neural-crest and CNS/sensory branches of the neural plate border network within a single regulatory unit.

### TE family identity determines co-option propensity and motif content

We next asked whether the four human-specific TE families investigated here (SVA-E, SVA-F, L1HS, LTR5Hs) differ in their propensity to be co-opted. Their relative representation among accessible TEs deviated markedly from their genomic abundance (chi-squared test, p = 3.03 × 10⁻^41^; **Figs. 3B, C**). SVA-F was enriched from 24.5% of expected to 41.2% of observed accessible TEs (Fisher’s exact test, p = 2.01 × 10⁻^9^; **Figs. 3B, C**), and LTR5Hs from 17.3% to 39.1% (p = 3.17 × 10⁻^17^; **Figs. 3B, C**). Conversely, SVA-E and L1HS were significantly depleted among accessible elements (p = 1.62 × 10⁻^43^, combined test; **Figs. 3B, C**). This depletion was especially notable for L1HS, which despite being the single most abundant human-specific TE family genome-wide (41.7% of all elements) constituted only 5.2% of accessible TEs. Furthermore, no accessible L1HS element contained any of the 15 enriched TFBMs, suggesting that L1HS elements are largely excluded from co-option in this cell population, in contrast to SVA-F and LTR5Hs.

We then asked whether TE family also determines the number of TFBMs harbored per accessible element. On average, accessible LTR5Hs elements contained more motifs per element than SVAs (5 vs. 3). Since SVAs contain an internal, length-variable VNTR domain, we considered whether this difference simply reflects element length rather than an intrinsic sequence property. Accessible SVAs had a shorter median length (180 bp) but were highly length-variable (IQR = 314 bp), whereas LTR5Hs elements were longer (median 964 bp) but length-conserved (IQR = 18 bp).

Notably, we observed family-specific bias extended to individual motifs. Among all motif-containing TEs, SVA-F accounted for 50.5% and LTR5Hs for 44.0%. The NEUROD2/ATOH1 motif was significantly biased toward SVA-F (68.9% of occurrences; Fisher’s exact test, p = 2.37 × 10⁻^7^; **Fig. 3D**), and the neural-crest-restricted SMAD4 motif was almost exclusively found within SVA elements, again predominantly SVA-F (88.9%; **Fig. 3D**). By contrast, the STAT3/PAX5 motif, occupying the neural-crest/CNS decision point itself, showed no significant family bias (chi-squared test, p = 0.78; **Figure 3D**), and the CRX motif was found exclusively within LTR5Hs elements. It is notable that the two motifs showing the strongest, most exclusive family restriction (SMAD4 to SVA; CRX to LTR5Hs) sit at opposite ends of the neural-crest/CNS functional spectrum, raising the possibility that SVA and LTR5Hs elements are biased toward different branches of the bipotent neural plate border program.

Motif count was strongly correlated with element length for SVAs (Pearson r = 0.77, p = 1.94 × 10⁻^24^) but not for LTR5Hs elements (**Fig. 4**; Supplementary File 8). Fitting a linear model of motif count against length using only SVA elements, and projecting LTR5Hs observations onto this model, produced a substantially larger residual error for LTR5Hs (RMSE = 10.1) than for SVA itself (RMSE = 3.19), indicating that the higher raw motif count observed in LTR5Hs is not a simple consequence of element length, and that SVA and LTR5Hs might achieve TFBM enrichment via distinct underlying sequence properties, likely reflecting the composite, internally repetitive structure of SVA elements versus the comparatively invariant, solo-LTR-derived structure of LTR5Hs.

**Figure 4.**
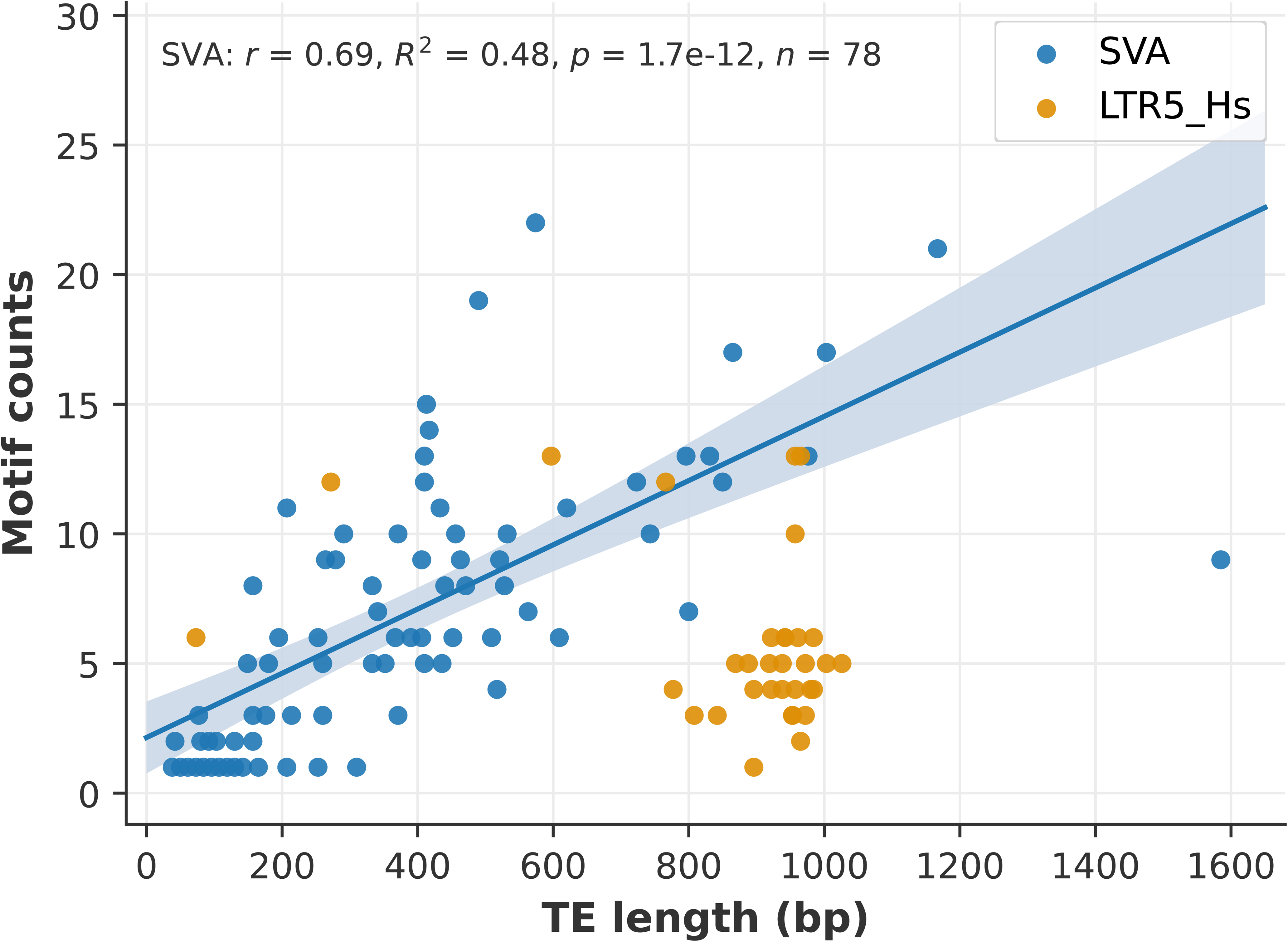
Motif content scales with element length in SVA but not LTR5Hs. Number of enriched transcription factor binding motifs per element plotted against element length, for SVA (n = 78) and LTR5_Hs (n = 33) elements. Line and shaded band show the linear regression and 95% confidence interval fitted to SVA elements (r = 0.69, R² = 0.48, P = 1.7 × 10⁻^12^). LTR5Hs elements show no relationship between length and motif count (r = −0.32, P = 0.07) and carry fewer motifs than the SVA trend predicts at equivalent length.

### Genes near accessible TEs are enriched for both craniofacial and CNS/sensory pathways

To delve into downstream biological processes potentially regulated by co-opted human-specific TE-CREs, we identified the unique genes located nearest to the 289 accessible TEs and performed Gene Ontology over-representation analysis. Ten gene sets were significantly enriched (p < 0.05; **Fig. 5A**; Supplementary File 9), and, consistent with the bipotent identity of the profiled population, these spanned both neural-crest-typical and CNS/sensory-typical categories rather than clustering exclusively around one of them.

**Figure 5.**
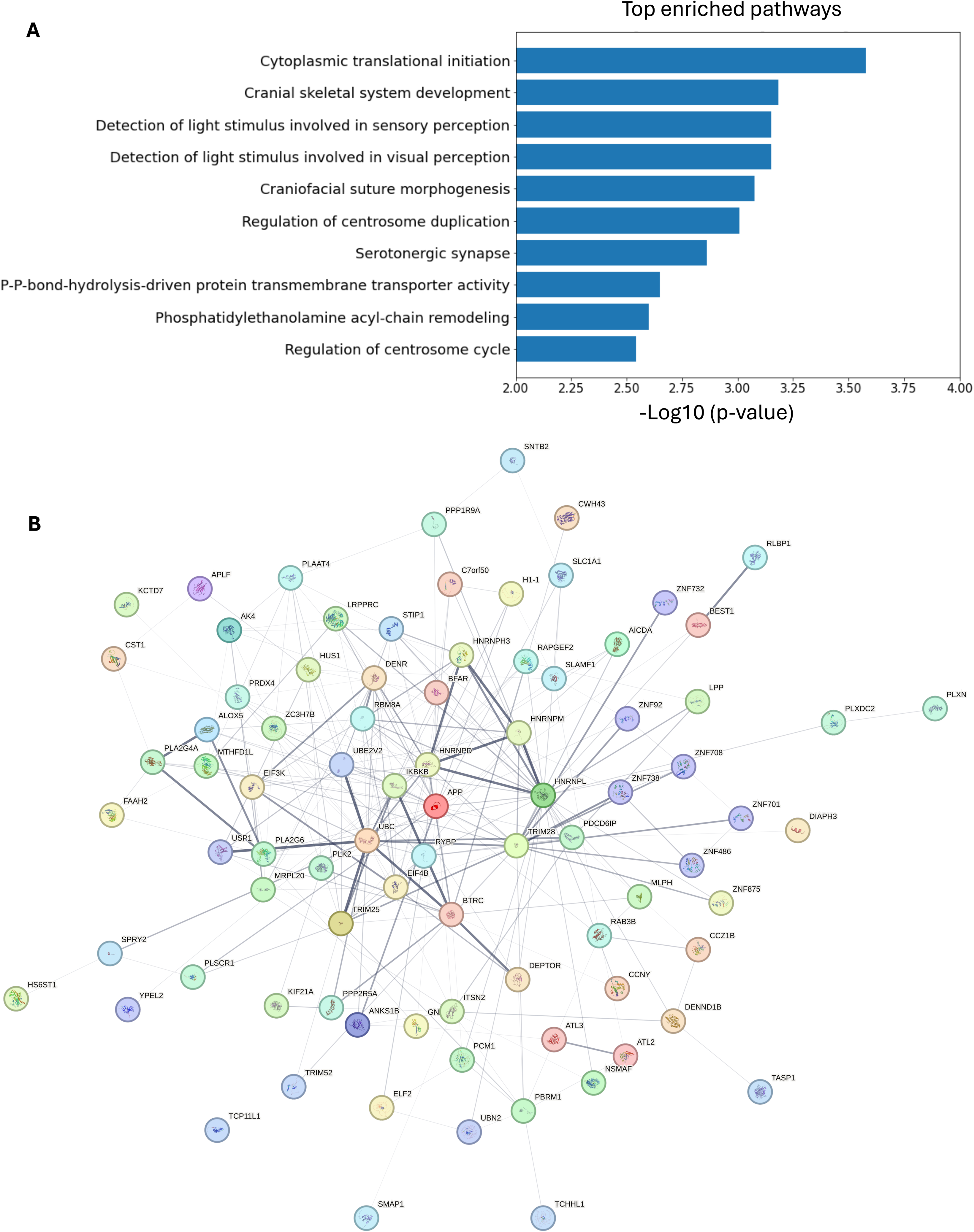
Genes near accessible TEs converge on craniofacial and sensory pathways and on TE-silencing machinery. (A) Top enriched pathways among genes associated with accessible human-specific TEs. (B) STRING network (v12.0, all sources active, minimum interaction score 0.150) of the same gene set. Edge thickness reflects interaction confidence. Node color indicates cluster and node fill shows protein structure.

Two enriched categories directly reflected neural-crest-derived craniofacial skeletal development (GO:1904888, GO:0097094), containing *FREM1*, typically mutated in metopic craniosynostosis, a congenital craniofacial suture defect (Vissers et al. 2011), and *INSIG1/2*, negative regulators of cholesterol homeostasis that modulate Wnt/BMP-dependent craniofacial osteogenic differentiation of NCC and whose loss reduces frontal bone formation in mouse knockouts (Suzuki et al. 2022). A further two categories were explicitly CNS/sensory: light-detection and visual perception (GO:0050908, GO:0050962), containing *BEST1*, a direct transcriptional target of CRX in retinal pigment epithelium, directly corroborating the CRX motif enrichment identified in our TFBM analysis above, and serotonergic synapse signaling, containing the serotonin receptor genes *HTR1B* and *HTR2C*. Serotonergic signaling is a purely neuronal/CNS pathway with no established role in neural crest mesenchymal differentiation. A further enriched category, centrosome duplication/cycle (GO:0046605, GO:0010824), relates to cell-cycle dynamics that have been linked in other systems to the differential migratory behavior of neural crest cells arising from different axial levels, and may similarly reflect the underlying cellular heterogeneity of the population.

To assess whether the proteins encoded by the 187 genes associated with accessible human-specific TEs show evidence of functional association, we employed STRING (Szklarczyk et al. 2023; 2025; FDR < 0.05; **Fig. 5B**). The majority of the proteins were connected within a single component, with UBC and TRIM28 as the principal hubs. It is worth noting that UBC is a recurrent hub in protein interaction networks due to the promiscuity of ubiquitin conjugation. On the other hand, the most coherent module comprised TRIM28 together with seven KRAB zinc-finger proteins (ZNF92, ZNF486, ZNF701, ZNF708, ZNF732, ZNF738 and ZNF875; **Fig. 5B**). A second module comprised heterogeneous nuclear ribonucleoproteins and associated RNA-binding factors (HNRNPL, HNRNPM, HNRNPD, HNRNPH3, RBM8A, DENR and EIF4B; **Fig. 5B**), and a third the phospholipase A2 enzymes with ALOX5 (PLA2G4A, PLA2G6, ALOX5; **Fig. 5B**). Together, the co-occurrence of craniofacial-skeletal and CNS/sensory/serotonergic enrichment within a single unstratified gene list is consistent with the expected molecular signature of a bipotent, pre-migratory NCC/CNS precursor population.

## Discussion

This study set out to characterize human-specific TE co-option within the bipotent, neural-plate-border-equivalent neuroectodermal population from which both the NCC and CNS/sensory lineages emerge. Consistent with this, the transcription factor binding motifs enriched within accessible TEs span the full functional spectrum of the neural plate border network, from CNS/sensory terminal differentiation factors (NEUROD2/ATOH1, CRX) through the neural-crest/CNS decision-point factor STAT3, to neural-crest-restricted differentiation factors (SMAD4). Moreover, the genes located nearest to accessible TEs are enriched for both neural-crest-typical (craniofacial skeletal) and CNS-typical (visual/light detection, serotonergic synapse) pathways within the same, non-stratified gene list. Taken together, these observations indicate that the TE-associated regulatory landscape captured here is a molecular readout of the developmental bipotency of the neuroectodermal aggregate model. A regulatory element active within a bipotent precursor population is positioned to influence the initial partitioning of cell fate between the neural crest and CNS/sensory lineages, rather than simply fine-tuning an already-committed neural crest program. Human-specific TE co-option at this stage could therefore, in principle, contribute not only to craniofacial evolutionary divergence but also to the parallel, and potentially coupled, evolutionary divergence of CNS and cranial sensory structures, an idea consistent with previous studies on the divergence between humans and chimpanzees in auditory sensitivity (Kojima 1990; Coleman and Ross 2004). The joint enrichment of ATOH1 (inner ear hair cell specification) and CRX (photoreceptor specification) motifs within our accessible TE set, alongside craniofacial skeletal pathway genes, is consistent with this possibility.

The pronounced family-specific co-option bias we observed, showing enrichment of SVA-F and LTR5Hs, and depletion of SVA-E and L1HS, and largely non-overlapping, length-independent motif signatures for SVA versus LTR5Hs, recapitulates family-specific biases previously reported in human pluripotent stem cells, hippocampal progenitors and migratory neural crest cells (Barnada et al., 2022; Patoori et al., 2022; Deelen et al. 2025), extending this pattern to the neural-plate-border-equivalent stage. The likely origin of this bias lies in family-specific “proto-motifs” retained within each TE family’s ancestral replication-associated sequence, which predispose different families toward acquiring different host TFBMs through subsequent substitution mutation (Bourque et al., 2008). The markedly greater sequence diversity of SVA elements, which are composite retrotransposons containing an internal VNTR domain, relative to the largely invariant, solo-LTR-derived LTR5Hs family, plausibly explains the broader and more length-dependent motif repertoire of SVA relative to LTR5Hs (Hughes & Coffin, 2004).

The near-mutually-exclusive family restriction of the neural-crest-specific SMAD4 motif to SVA elements and the CNS-specific CRX motif to LTR5Hs elements raises the possibility that these two TE families are biased towards opposite poles of the neural-crest/CNS regulatory spectrum. However, because our data derive from bulk chromatin accessibility profiling of a mixed, unresolved population, we cannot currently determine whether this apparent family/lineage association reflects two overlapping sets of TEs simultaneously accessible within single bipotent cells, later resolved differentially as each cell commits to one fate or the other, or two distinct subpopulations of cells within the aggregates, each already biased toward one fate and each preferentially using one TE family.

From a clinical and evolutionary perspective, our data suggest that human-specific TE co-option at this bipotent developmental stage could contribute to the pathology of a broader range of conditions than craniofacial birth defects alone. Since the majority of the SVA and LTR5Hs elements investigated here remain transcriptionally and (in the case of SVA) transpositionally active, polymorphic TE insertions with cis-regulatory consequences could plausibly contribute both to craniofacial malformation and, through the parallel CNS/sensory-associated regulatory signature we describe, to neurodevelopmental or neuropsychiatric phenotypes. The enrichment of serotonergic signaling pathway genes is particularly notable in this context, given that polymorphisms in serotonin receptor genes, including *HTR1B* and *HTR2C* identified within our own gene list, have previously been associated with schizophrenia, anxiety and suicidal behaviour (Serretti et al. 2007; Xia et al. 2018), and given that SVA and LTR5Hs elements are themselves highly polymorphic across human populations.

The recovery of a TRIM28–KZFP module from a gene set defined by accessible human-specific transposable elements is notable, because this is the machinery that silences those elements. KRAB zinc-finger proteins bind TE families in a sequence-specific manner and recruit TRIM28 (KAP1), which in turn recruits SETDB1 to deposit H3K9me3 and nucleate heterochromatin (Schultz et al., 2002; Rowe et al., 2010; Turelli et al., 2014). This relationship has been described as an evolutionary arms race, in which KZFP genes have expanded in primates in parallel with the retroelement families they target, and individual KZFPs show binding specificity for particular subfamilies (Thomas and Schneider, 2011; Jacobs et al., 2014; Imbeault et al., 2017; Helleboid et al., 2019). Our observation that genes proximal to accessible human-specific TEs are enriched for members of this module could have two interpretations. The first is regulatory feedback, in which de-repression of a TE family exposes elements that themselves regulate expression of their own silencing machinery, a configuration that would confer self-limiting control over TE activity. The second, and more likely given that our data are correlative, is that KZFP genes are recovered because they are physically embedded in TE-rich genomic territory: KZFP genes are organized in large tandem clusters, predominantly on chromosome 19, that are among the most repeat-dense regions of the human genome (Huntley et al., 2006; Thomas and Schneider, 2011), and their proximity to accessible elements may therefore reflect genomic context rather than functional regulation.

## Materials and Methods

### iPSC culture

Human iPSCs (line SV20, University of Pennsylvania) were maintained as monolayer cultures on Geltrex™-coated 6-well plates in feeder-free, serum-free mTeSR™ Plus medium (StemCell Technologies) supplemented with 1% penicillin-streptomycin, at 37°C, 5% CO₂, 20% O₂. Cells were passaged as small clumps with 0.5 mM EDTA every 3–4 days at ∼80% confluency (split ratio ∼1:6).

### iPSC differentiation to neuroectodermal aggregates

To detach iPSC colonies, cultures were treated with accutase. Detached colonies were then broken into clusters of approximately 100–200 cells and transferred to NCC differentiation medium, consisting of Neurobasal and D-MEM/F-12 media (Invitrogen) mixed 1:1 and supplemented with B-27 with Vitamin A at 0.5× (from 50× stock, Invitrogen), N-2 at 0.5× (from 100× stock, Invitrogen), bFGF at 20 ng/ml (Peprotech), EGF at 20 ng/ml (Sigma-Aldrich), bovine insulin at 5 µg/ml (Sigma-Aldrich) and Glutamax-I at 1× (from 100× stock, Invitrogen). Cultures were fed every second day, and aggregates were harvested on day 5 of differentiation.

### ATAC-seq library preparation and data analysis

ATAC-seq (Buenrostro et al. 2013) libraries were prepared from 50,000 day-5 aggregate cells in two technical replicates using Active Motif’s ATAC-Seq Kit. Fragment size was assessed by TapeStation. Libraries were sequenced on an Illumina NextSeq 550 (150 bp paired-end) at the Genomics Facility of the Wistar Institute (Philadelphia, PA). Adaptors were trimmed with Trim Galore! (Krueger et al. 2021) and reads mapped to hg19 with BWA-MEM (Li 2013); reads with MAPQ > 10 were retained and PCR duplicates removed with SAMtools (Danacek et al. 2021). Peaks were called with MACS2 (Zhang et al. 2008; 5% FDR).

### Accessible TE identification and motif analysis

Human-specific TE coordinates (UCSC RepeatMasker, hg19) were intersected with ATAC-seq accessibility using deepTools (Ramirez et al. 2016; computeMatrix/plotHeatmap, ±1 kb around each TE, k-means clustering). Distance from each TE to the nearest annotated TSS was calculated with BEDTools closest (v2.30.0; Quinlan et al. 2010). Motif enrichment within accessible TE sequences was performed with MEME-ChIP (Machanick and Bailey 2011; default parameters, motifs ≥ 6 nt), and motif locations within TEs were mapped with MEME-FIMO (E < 1 × 10⁻^5^). For the motif analysis, non-accessible human-specific TEs were used as control group.

### RNA library preparation

RNA was extracted from frozen day-0 (iPSC) and day-5 pellets using the Monarch® Total RNA Miniprep Kit and quantified by NanoDrop. RNA integrity was assessed prior to library preparation, and only samples with a RIN above 8 were taken forward. For each library, 1 µg of total RNA was used as input. Poly(A) selection was carried out with the NEBNext Poly(A) mRNA Magnetic Isolation Module (NEB, E7490), followed by library preparation with the NEBNext Ultra II Directional RNA Library Prep Kit for Illumina (NEB, E7760). Libraries were sequenced by The Wistar Institute Sequencing Facility (Philadelphia, PA).

### RNA-seq quantification and gene-level summarization

Transcript-level abundances were estimated with Kallisto (Bray et al. 2016) against an index built from the Ensembl release 96 human cDNA set (GRCh38; 188,753 transcripts). Transcript estimates were summarized to genes following the tximport procedure, implemented in Python: estimated counts were summed per gene, and a per-gene effective length was computed as the abundance-weighted mean of constituent transcript effective lengths. Two matrices were generated: raw summed counts with the accompanying average-transcript-length offset matrix, and a length-scaled TPM matrix (countsFromAbundance = “lengthScaledTPM”).

### Differential expression

Differential expression between day5 (n = 3) and day-0 cells (n = 2) was tested with DESeq2 (Love et al. 2014). Gene-wise dispersions were estimated by maximum likelihood, fitted to a parametric mean–dispersion trend, and shrunk towards the trend by empirical Bayes. Differential expression was tested under a negative binomial generalized linear model with the design ∼ condition, using the Wald test on the day-5 versus day-0 coefficient. P-values were adjusted by the Benjamini–Hochberg procedure following DESeq2 independent filtering on mean normalized count. After filtering, 17,852 genes were tested and genes with adjusted P < 0.05 were considered differentially expressed.

### Over-representation analysis

Upregulated genes (adjusted P < 0.05) were tested for over-representation against gene set collections with the WebGestalt API (Elizarraras et al. 2024), and in particular: Gene Ontology Biological Process (non-redundant), KEGG, Reactome and WikiPathways. Over-representation was assessed by hypergeometric test, with the set of genes retained after DESeq2 independent filtering as the background. P-values were adjusted for multiple testing by the Benjamini– Hochberg correction within each database, and terms with FDR < 0.05 were considered enriched.

### Distance to TSS and gene density

For each element, distance to the nearest TSS was computed as the absolute distance to the closest annotated TSS on the same chromosome, and gene density was computed as the number of TSSs within a fixed window. Accessible and non-accessible elements were compared by two-sided Mann–Whitney U test, with Cliff’s delta reported as a non-parametric effect size.

### Expression of neighboring genes

Each element was assigned to its nearest quantified gene, and where several elements shared a nearest gene, only the closest element was retained to avoid pseudoreplication. Expression of genes neighboring accessible versus non-accessible elements was compared on log2-transformed day-5 values, using both length-scaled TPM and DESeq2 median-of-ratios normalized counts, by two-sided Mann–Whitney U test with Cliff’s delta as effect size.

Because accessible elements differ from non-accessible elements in both subfamily composition and distance to TSS, comparisons were repeated against matched control sets. For each accessible element, a non-accessible element was sampled without replacement from the same subfamily and the same distance-to-TSS bin. Matching was repeated over 500 resamples, and the median Cliff’s delta, median P-value and proportion of resamples with P < 0.05 are reported. To establish the false-positive rate expected from sampling variation alone, 1,000 further comparisons were performed against unmatched random draws from the non-accessible pool of equal size to the accessible set. An additional window-based analysis considered all quantified genes whose TSS fell within specific distances of an element, rather than the single nearest gene, and compared the union of genes near accessible versus near matched non-accessible elements by the same tests.

### Protein interaction network

Genes associated with accessible human-specific elements (n = 87) were queried against STRING v12.0 (Homo sapiens), with all interaction sources active and a minimum required interaction score of 0.150. Clusters were defined by k-means, with 3 clusters. Two entries (MIR7-1, SUZ12P1) were not mapped, as STRING is restricted to protein-coding genes.

### Statistical reporting

All tests were two-sided. Non-parametric tests were used throughout for comparisons of expression and genomic distance. Effect sizes are reported as Cliff’s delta, which is bounded by −1 and +1 and independent of the scale of the underlying measurements. Multiple testing was controlled by the Benjamini–Hochberg correction at 5% FDR. Analyses were performed in Python 3.12 using pandas, NumPy, SciPy.

### Data availability statement

RNA-seq and ATAC-seq data are available the Gene Expression Omnibus (GEO) under accession code GEO: GSE293821 (RNA-seq) and GEO: GSE347017 (ATAC-seq).

## Acknowledgements

We thank the members of the Trizzino group for insightful discussions on the data.

## Author contributions

MN performed the experiments. ZM contributed to the experiments. MN and MT analyzed the data and drafted the first draft. MT designed and supervised the project, revised and prepared the manuscript for submission.

## References

Bajpai, R., D. A. Chen, A. Rada-Iglesias, J. Zhang, Y. Xiong, et al., 2010 CHD7 cooperates with PBAF to control multipotent neural crest formation. Nature 463: 958–962. 10.1038/nature08733

Barnada, S. M., A. Isopi, D. Tejada-Martinez, C. Goubert, S. Patoori, et al., 2022 Genomic features underlie the co-option of SVA transposons as cis-regulatory elements in human pluripotent stem cells. PLoS Genet. 18: e1010225. 10.1371/journal.pgen.1010225

Bermingham, N. A., B. A. Hassan, S. D. Price, M. A. Vollrath, N. Ben-Arie, et al., 1999 Math1: an essential gene for the generation of inner ear hair cells. Science 284: 1837–1841. 10.1126/science.284.5421.1837

Bourque, G., B. Leong, V. B. Vega, X. Chen, Y. L. Lee, et al., 2008 Evolution of the mammalian transcription factor binding repertoire via transposable elements. Genome Res. 18: 1752–1762. 10.1101/gr.080663.108

Bray, N. L., H. Pimentel, P. Melsted, and L. Pachter, 2016 Near-optimal probabilistic RNA-seq quantification. Nat. Biotechnol. 34: 525–527. 10.1038/nbt.3519

Bronner, M. E., and M. Simões-Costa, 2016 The neural crest migrating into the twenty-first century. Curr. Top. Dev. Biol. 116: 115–134. 10.1016/bs.ctdb.2015.12.003

Buenrostro, J. D., P. G. Giresi, L. C. Zaba, H. Y. Chang, and W. J. Greenleaf, 2013 Transposition of native chromatin for fast and sensitive epigenomic profiling of open chromatin, DNA-binding proteins and nucleosome position. Nat. Methods 10: 1213–1218. 10.1038/nmeth.2688

Chuong, E. B., M. A. K. Rumi, M. J. Soares, and J. C. Baker, 2013 Endogenous retroviruses function as species-specific enhancer elements in the placenta. Nat. Genet. 45: 325–329. 10.1038/ng.2553

Chuong, E. B., N. C. Elde, and C. Feschotte, 2016 Regulatory evolution of innate immunity through co-option of endogenous retroviruses. Science 351: 1083–1087. 10.1126/science.aad5497

Coleman, M. N., and C. F. Ross, 2004 Primate auditory diversity and its influence on hearing performance. Anat. Rec. A 281: 1123–1137. 10.1002/ar.a.20118

Danecek, P., J. K. Bonfield, J. Liddle, J. Marshall, V. Ohan, et al., 2021 Twelve years of SAMtools and BCFtools. GigaScience 10: giab008. 10.1093/gigascience/giab008

Deelen, L., Z. H. Mitchell, M. Demurtas, A. Koulle, B. Garcia Del Valle, et al., 2025 Hominoid-specific transposable elements reshaped neural crest migration in craniofacial development. Mol. Syst. Biol. 21: 1731–1747. 10.1038/s44320-025-00151-z

Demurtas, M., S. M. Barnada, E. van Domselaar, Z. H. Mitchell, L. Deelen, et al., 2025 Neural crest induction requires SALL4-mediated BAF recruitment to lineage specific enhancers. Development 152: dev205248. 10.1242/dev.205248

Elizarraras, J. M., Y. Liao, Z. Shi, Q. Zhu, A. R. Pico, et al., 2024 WebGestalt 2024: faster gene set analysis and new support for metabolomics and multi-omics. Nucleic Acids Res. 52: W415– W421. 10.1093/nar/gkae456

Frost, J. M., S. M. Amante, H. Okae, E. M. Jones, B. Ashley, et al., 2023 Regulation of human trophoblast gene expression by endogenous retroviruses. Nat. Struct. Mol. Biol. 30: 527–538. 10.1038/s41594-023-00960-6

Fueyo, R., J. Judd, C. Feschotte, and J. Wysocka, 2022 Roles of transposable elements in the regulation of mammalian transcription. Nat. Rev. Mol. Cell Biol. 23: 481–497. 10.1038/s41580-022-00457-y

Fueyo, R., S. Wang, O. J. Crocker, T. Swigut, H. Nakauchi, et al., 2025 A human-specific regulatory mechanism revealed in a pre-implantation model. Nature 647: 238–247. 10.1038/s41586-025-09571-1

Furukawa, T., E. M. Morrow, T. Li, F. C. Davis, and C. L. Cepko, 1999 Retinopathy and attenuated circadian entrainment in Crx-deficient mice. Nat. Genet. 23: 466–470. 10.1038/70591

Grant, C. E., T. L. Bailey, and W. S. Noble, 2011 FIMO: scanning for occurrences of a given motif. Bioinformatics 27: 1017–1018. 10.1093/bioinformatics/btr064

Groves, A. K., and C. LaBonne, 2014 Setting appropriate boundaries: fate, patterning and competence at the neural plate border. Dev. Biol. 389: 2–12. 10.1016/j.ydbio.2013.11.027

Hancks, D. C., and H. H. Kazazian, Jr., 2010 SVA retrotransposons: evolution and genetic instability. Semin. Cancer Biol. 20: 234–245. 10.1016/j.semcancer.2010.04.001

Helleboid, P.-Y., M. Heusel, J. Duc, C. Piot, C. W. Thorball, et al., 2019 The interactome of KRAB zinc finger proteins reveals the evolutionary history of their functional diversification. EMBO J. 38: e101220. 10.15252/embj.2018101220

Hidalgo-Sánchez, M., A. Andreu-Cervera, S. Villa-Carballar, and D. Echevarria, 2022 An update on the molecular mechanism of the vertebrate isthmic organizer development in the context of the neuromeric model. Front. Neuroanat. 16: 826976. 10.3389/fnana.2022.826976

Hughes, J. F., and J. M. Coffin, 2004 Human endogenous retrovirus K solo-LTR formation and insertional polymorphisms: implications for human and viral evolution. Proc. Natl. Acad. Sci. USA 101: 1668–1672. 10.1073/pnas.0307885100

Huntley, S., D. M. Baggott, A. T. Hamilton, M. Tran-Gyamfi, S. Yang, et al., 2006 A comprehensive catalog of human KRAB-associated zinc finger genes: insights into the evolutionary history of a large family of transcriptional repressors. Genome Res. 16: 669–677. 10.1101/gr.4842106

Imbeault, M., P.-Y. Helleboid, and D. Trono, 2017 KRAB zinc-finger proteins contribute to the evolution of gene regulatory networks. Nature 543: 550–554. 10.1038/nature21683

Jacobs, F. M. J., D. Greenberg, N. Nguyen, M. Haeussler, A. D. Ewing, et al., 2014 An evolutionary arms race between KRAB zinc-finger genes ZNF91/93 and SVA/L1 retrotransposons. Nature 516: 242–245. 10.1038/nature13760

Jacques, P.-É., J. Jeyakani, and G. Bourque, 2013 The majority of primate-specific regulatory sequences are derived from transposable elements. PLoS Genet. 9: e1003504. 10.1371/journal.pgen.1003504

Keuls, R. A., Y. S. Oh, I. Patel, and R. J. Parchem, 2023 Post-transcriptional regulation in cranial neural crest cells expands developmental potential. Proc. Natl. Acad. Sci. USA 120: e2212578120. 10.1073/pnas.2212578120

Ko, S. O., I. H. Chung, X. Xu, S. Oka, H. Zhao, et al., 2007 Smad4 is required to regulate the fate of cranial neural crest cells. Dev. Biol. 312: 435–447. 10.1016/j.ydbio.2007.09.050

Kojima, S., 1990 Comparison of auditory functions in the chimpanzee and human. Folia Primatol. 55: 62–72. 10.1159/000156501

Krueger, F., F. James, P. Ewels, E. Afyounian, and B. Schuster-Böckler, 2021 FelixKrueger/TrimGalore: v0.6.7. Zenodo. 10.5281/zenodo.5127899

Kunarso, G., N.-Y. Chia, J. Jeyakani, C. Hwang, X. Lu, et al., 2010 Transposable elements have rewired the core regulatory network of human embryonic stem cells. Nat. Genet. 42: 631–634. 10.1038/ng.600

Lencer, E., R. Prekeris, and K. B. Artinger, 2021 Single-cell RNA analysis identifies pre-migratory neural crest cells expressing markers of differentiated derivatives. eLife 10: e66078. 10.7554/eLife.66078

Li, H., 2013 Aligning sequence reads, clone sequences and assembly contigs with BWA-MEM. arXiv:1303.3997.

Li, J., X. Huang, X. Xu, J. Mayo, P. Bringas, Jr.,et al ., 2011 SMAD4-mediated WNT signaling controls the fate of cranial neural crest cells during tooth morphogenesis. Development 138: 1977–1989. 10.1242/dev.061341

Long, H. K., M. Osterwalder, I. C. Welsh, K. Hansen, J. O. J. Davies, et al., 2020 Loss of extreme long-range enhancers in human neural crest drives a craniofacial disorder. Cell Stem Cell 27: 765–783.e14. 10.1016/j.stem.2020.09.001

Love, M. I., W. Huber, and S. Anders, 2014 Moderated estimation of fold change and dispersion for RNA-seq data with DESeq2. Genome Biol. 15: 550. 10.1186/s13059-014-0550-8

Lumb, R., S. Buckberry, G. Secker, D. Lawrence, and Q. Schwarz, 2017 Transcriptome profiling reveals expression signatures of cranial neural crest cells arising from different axial levels. BMC Dev. Biol. 17: 20. 10.1186/s12861-017-0161-1

Lynch, V. J., R. D. Leclerc, G. May, and G. P. Wagner, 2011 Transposon-mediated rewiring of gene regulatory networks contributed to the evolution of pregnancy in mammals. Nat. Genet. 43: 1154–1159. 10.1038/ng.917

Lynch, V. J., M. C. Nnamani, A. Kapusta, K. Brayer, S. L. Plaza, et al., 2015 Ancient transposable elements transformed the uterine regulatory landscape and transcriptome during the evolution of mammalian pregnancy. Cell Rep. 10: 551–561. 10.1016/j.celrep.2014.12.052

Machanick, P., and T. L. Bailey, 2011 MEME-ChIP: motif analysis of large DNA datasets. Bioinformatics 27: 1696–1697. 10.1093/bioinformatics/btr189

Markljung, E., L. Jiang, J. D. Jaffe, T. S. Mikkelsen, O. Wallerman, et al., 2009 ZBED6, a novel transcription factor derived from a domesticated DNA transposon regulates IGF2 expression and muscle growth. PLoS Biol. 7: e1000256. 10.1371/journal.pbio.1000256

Martin, M., 2011 Cutadapt removes adapter sequences from high-throughput sequencing reads. EMBnet J. 17: 10–12. 10.14806/ej.17.1.200

Mayor, R., and E. Theveneau, 2013 The neural crest. Development 140: 2247–2251. 10.1242/dev.091751

Mitchell, Z. H., J. den Hoed, W. Claassen, M. Demurtas, L. Deelen, et al., 2025 The NuRD component CHD3 promotes BMP signalling during cranial neural crest cell specification. EMBO Rep. 26: 4723–4741. 10.1038/s44319-025-00555-w

Nichane, M., X. Ren, and E. J. Bellefroid, 2010 Self-regulation of Stat3 activity coordinates cell-cycle progression and neural crest specification. EMBO J. 29: 55–67. 10.1038/emboj.2009.313

Ozga, E., K. M. Milto, M. Demurtas, L. E. Bates, G. Grimes, et al., 2026 Array-CNCC: precise aggregation and arrayed plating facilitate quantitative phenotyping of human cranial neural crest cells and craniofacial disease modelling. bioRxiv 2026.01.18.696654. 10.1101/2026.01.18.696654

Patoori, S., S. M. Barnada, C. Large, J. I. Murray, and M. Trizzino, 2022 Young transposable elements rewired gene regulatory networks in human and chimpanzee hippocampal intermediate progenitors. Development 149: dev200413. 10.1242/dev.200413

Pontis, J., E. Planet, S. Offner, P. Turelli, J. Duc, et al., 2019 Hominoid-specific transposable elements and KZFPs facilitate human embryonic genome activation and control transcription in naive human ESCs. Cell Stem Cell 24: 724–735.e5. 10.1016/j.stem.2019.03.012

Prescott, S. L., R. Srinivasan, M. C. Marchetto, I. Grishina, I. Narvaiza, et al., 2015 Enhancer divergence and cis-regulatory evolution in the human and chimp neural crest. Cell 163: 68–83. 10.1016/j.cell.2015.08.036

Quinlan, A. R., and I. M. Hall, 2010 BEDTools: a flexible suite of utilities for comparing genomic features. Bioinformatics 26: 41–842. 10.1093/bioinformatics/btq033

Ramírez, F., D. P. Ryan, B. Grüning, V. Bhardwaj, F. Kilpert, et al., 2016 deepTools2: a next generation web server for deep-sequencing data analysis. Nucleic Acids Res. 44: W160–W165. 10.1093/nar/gkw257

Rowe, H. M., J. Jakobsson, D. Mesnard, J. Rougemont, S. Reynard, et al., 2010 KAP1 controls endogenous retroviruses in embryonic stem cells. Nature 463: 237–240. 10.1038/nature08674

Sasaki, T., A. Nishihara, M. Hirakawa, K. Fujimura, M. Santangelo, et al., 2008 Possible involvement of SINEs in mammalian-specific brain formation. Proc. Natl. Acad. Sci. USA 105: 4220–4225. 10.1073/pnas.0709398105

Schmidt, D., P. C. Schwalie, M. D. Wilson, B. Ballester, A. Gonçalves, et al., 2012 Waves of retrotransposon expansion remodel genome organization and CTCF binding in multiple mammalian lineages. Cell 148: 335–348. 10.1016/j.cell.2011.11.058

Schultz, D. C., K. Ayyanathan, D. Negorev, G. G. Maul, and F. J. Rauscher, 3rd, 2002 SETDB1: a novel KAP-1-associated histone H3, lysine 9-specific methyltransferase that contributes to HP1-mediated silencing of euchromatic genes by KRAB zinc-finger proteins. Genes Dev. 16: 919–932. 10.1101/gad.973302

Serretti, A., L. Mandelli, I. Giegling, B. Schneider, A. M. Hartmann, et al., 2007 HTR2C and HTR1A gene variants in German and Italian suicide attempters and completers. Am. J. Med. Genet. B 144: 291–299. 10.1002/ajmg.b.30432

Simões-Costa, M., and M. E. Bronner, 2015 Establishing neural crest identity: a gene regulatory recipe. Development 142: 242–257. 10.1242/dev.105445

Simões-Costa, M., J. Tan-Cabugao, I. Antoshechkin, T. Sauka-Spengler, and M. E. Bronner, 2014 Transcriptome analysis reveals novel players in the cranial neural crest gene regulatory network. Genome Res. 24: 281–290. 10.1101/gr.161182.113

Sundaram, V., Y. Cheng, Z. Ma, D. Li, X. Xing, et al., 2014 Widespread contribution of transposable elements to the innovation of gene regulatory networks. Genome Res. 24: 1963– 1976. 10.1101/gr.168872.113

Suzuki, A., K. Ogata, H. Yoshioka, J. Shim, C. A. Wassif, et al., 2020 Disruption of Dhcr7 and Insig1/2 in cholesterol metabolism causes defects in bone formation and homeostasis through primary cilium formation. Bone Res. 8: 1. 10.1038/s41413-019-0078-3

Szklarczyk, D., R. Kirsch, M. Koutrouli, K. Nastou, F. Mehryary, et al., 2023 The STRING database in 2023: protein–protein association networks and functional enrichment analyses for any sequenced genome of interest. Nucleic Acids Res. 51: D638–D646. 10.1093/nar/gkac1000

Szklarczyk, D., K. Nastou, M. Koutrouli, R. Kirsch, F. Mehryary, et al., 2025 The STRING database in 2025: protein networks with directionality of regulation. Nucleic Acids Res. 53: D730– D737. 10.1093/nar/gkae1113

Thomas, J. H., and S. Schneider, 2011 Coevolution of retroelements and tandem zinc finger genes. Genome Res. 21: 1800–1812. 10.1101/gr.121749.111

Trizzino, M., Y. Park, M. Holsbach-Beltrame, K. Aracena, K. Mika, et al., 2017 Transposable elements are the primary source of novelty in primate gene regulation. Genome Res. 27: 1623– 1633. 10.1101/gr.218149.116

Trizzino, M., A. Kapusta, and C. D. Brown, 2018 Transposable elements generate regulatory novelty in a tissue-specific fashion. BMC Genomics 19: 468. 10.1186/s12864-018-4850-3

Turelli, P., N. Castro-Diaz, F. Marzetta, A. Kapopoulou, C. Raclot, et al., 2014 Interplay of TRIM28 and DNA methylation in controlling human endogenous retroelements. Genome Res. 24: 1260–1270. 10.1101/gr.172833.114

Urbánek, P., I. Fetka, M. H. Meisler, and M. Busslinger, 1997 Cooperation of Pax2 and Pax5 in midbrain and cerebellum development. Proc. Natl. Acad. Sci. USA 94: 5703–5708. 10.1073/pnas.94.11.5703

Vissers, L. E. L. M., T. C. Cox, A. M. Maga, K. M. Short, F. Wiradjaja, et al., 2011 Heterozygous mutations of FREM1 are associated with an increased risk of isolated metopic craniosynostosis in humans and mice. PLoS Genet. 7: e1002278. 10.1371/journal.pgen.1002278

Wang, H., J. Xing, D. Grover, D. J. Hedges, K. Han, et al., 2005 SVA elements: a hominid-specific retroposon family. J. Mol. Biol. 354: 994–1007. 10.1016/j.jmb.2005.09.085

Wang, T., J. Zeng, C. B. Lowe, R. G. Sellers, S. R. Salama, et al., 2007 Species-specific endogenous retroviruses shape the transcriptional network of the human tumor suppressor protein p53. Proc. Natl. Acad. Sci. USA 104: 18613–18618. 10.1073/pnas.0703637104

Xia, X., M. Ding, J. Xuan, J. Xing, H. Pang, et al., 2018 Polymorphisms in the human serotonin receptor 1B (HTR1B) gene are associated with schizophrenia: a case control study. BMC Psychiatry 18: 9. 10.1186/s12888-017-1587-5

Zhang, Y., T. Liu, C. A. Meyer, J. Eeckhoute, D. S. Johnson, et al., 2008 Model-based analysis of ChIP-Seq (MACS). Genome Biol. 9: R137. 10.1186/gb-2008-9-9-r137

Zheng, Y., and S. Chen, 2024 Transcriptional precision in photoreceptor development and diseases – lessons from 25 years of CRX research. Front. Cell. Neurosci. 18: 1347436. 10.3389/fncel.2024.1347436

